# Introduced populations of ragweed show as much evolutionary potential as native populations

**DOI:** 10.1101/305540

**Authors:** Brechann V. McGoey, John R. Stinchcombe

## Abstract

Invasive species are a global economic and ecological problem. They also offer an opportunity to understand evolutionary processes in a colonizing context. The impacts of evolutionary factors, such as genetic variation, on the invasion process are increasingly appreciated but there remain gaps in the empirical literature. The adaptive potential of populations can be quantified using genetic variance-covariance matrices **(G)**, which encapsulate the heritable genetic variance in a population. Here, we use a multivariate, Bayesian approach to assess the adaptive potential of introduced populations of ragweed, *Ambrosia artemisiifolia*, a serious allergen and agricultural weed. We compared several aspects of genetic architecture and the structure of G matrices between three native and three introduced populations, based on data collected in the field in a common garden experiment. We find moderate differences in the quantitative genetic architecture among populations, but we do not find that introduced populations suffer from a limited adaptive potential compared to native populations. Ragweed has an annual life history, is an obligate outcrosser, and produces billions of seeds and pollen grains per. These characteristics, combined with the significant additive genetic variance documented here, suggest ragweed will be able to respond quickly to selection pressures in both its native and introduced ranges.

## Introduction

Anthropogenic change is altering the environment for species across the globe. One important way humans are changing the ecological landscape is through the accidental or intentional movement of organisms into novel locations. The ecological and economic impacts of alien plants continue to be immense (Sakai et al. 2001), generating an impetus to better understand how and why certain plant populations become invasive. Whether and how invasive species evolve in their new range is key to understanding their establishment and success (Colautti and Barrett 2013), yet we have a weak understanding of the evolutionary potential of size, performance, and life history traits in introduced species. Here we use a quantitative genetic approach to compare multivariate evolutionary potential between introduced and native populations of the prolific invader *Ambrosia artemisiifolia* (common ragweed).

Most research on species invasions has focused on possible ecological explanations and consequences, while the evolutionary determinants and outcomes have only recently been emphasized, and remain less well understood (Bacigalupe 2009). Recent evidence suggests that evolutionary responses can be more important in determining invasion success and spread than traditional ecological explanations (e.g., local adaptation accounting for more fitness effects than enemy release and allocation to competitive ability (EICA) (Colautti and Barrett 2013). Whereas there has long been an emphasis on individual level traits, it is increasingly recognized that population-level factors such as genetic variation, will have major impacts on the ability of a species to establish in a new environment and to respond to natural selection (Bacigalupe 2009; Crawford and Whitney 2010). Almost all traits that are likely to be under selection in a new environment are quantitative (Dlugosch and Parker 2008a), therefore characterizing the quantitative genetic variation of invasive populations is critical for understanding invasion success.

Despite the importance of heritable genetic variation for the ability of a population to respond to selection, there is a dearth of studies on invasive species from a quantitative genetics perspective (Bacigalupe 2009) and neither theory nor logic offer straightforward *a priori* predictions. While there is often an assumption that all introduced populations will suffer from founder effects, an initial bottleneck can be mitigated by multiple introductions (Roman and Darling 2007). Patterns of neutral genetic variation are unlikely to be helpful, as they are often uncorrelated with heritable quantitative variation (Reed and Frankham 2001; Mittell et al. 2015). Quantitative variation will be less impacted by losses of rare alleles (Dlugosch and Parker 2008a) and some theory and empirical studies suggest epistatic and dominance variance can actually be converted to additive variance (Bryant et al. 1986; Goodnight 1988; Willis and Orr 1993; Cheverud and Routman 1996), but see (Barton and Turelli 2004; Turelli and Barton 2006). Many studies of introduced species also suffer from insufficient sampling replication within the introduced and native ranges (Colautti et al. 2009). Ultimately, understanding an invasive species’ capacity to respond to selection and evolve is an empirical question which requires directly comparing additive genetic variation (V_A_) and covariation (**G**) in multiple introduced and native populations.

A well-established literature on variation in single-traits has uncovered genetic variance in almost all of them (Lynch and Walsh), which can falsely lead to the assumption that limited genetic variance is not a significant barrier to adaptive evolution (Blows and Hoffmann 2005). Even if there is additive genetic variance for a univariate trait, there can still be genetic constraints on adaptation due to covariances with other traits (Lande and Arnold 1983; McGuigan 2006; Agrawal and Stinchcombe 2009). The way a population moves across an adaptive landscape will be dictated by the available variance in many traits and the covariances between them (i.e., the genetic covariance matrix, **G**). A multivariate perspective, incorporating the impacts of multiple traits and their genetic correlations, is necessary for a comprehensive understanding of the available genetic variation in a population (Blows and Hoffmann 2005; Blows 2007; Walsh and Blows 2009).

The **G** matrix summarizes the available genetic variances and covariances, and offers an integrated view of quantitative genetic variation which allows for the estimation of constraints (Lande 1979; Blows 2007). Since it includes limitations caused by relationships among traits, it can uncover constraints on adaptation, even in cases where all the traits themselves have genetic variation (Dickerson 1955; Blows 2007). The **G** matrix will dictate the speed and direction of a population’s response to selection (Steppan et al. 2002). Understanding evolutionary potential in invasive species, therefore, requires a comparison of **G** matrices between invasive and native populations to determine whether invasive species face genetic constraints and will have reduced phenotypic evolutionary potential.

In this study, we examine how quantitative genetic architecture varies between introduced and native populations of the prolific invader *Ambrosia artemisiifolia* (ragweed). We focus on size and phenology traits that are likely to be under selection. Specifically, we ask, 1) How do native and introduced populations differ in their mean phenotypes, genetic variances, and heritabilities in key phenotypic traits? 2) Are there correlations between traits that could accelerate or constrain adaptation? 3) Is there a divergence between continents in the **G** matrices of populations? 4) How would native and invasive populations differ in their responses to selection based on their genetic (co)variances?

## Methods

### Study species

*Ambrosia artemisiifolia* is an annual herb (Bassett and Crompton 1975) and is self-incompatible (Friedman and Barrett 2008; Li et al. 2012). Preferring open habitats, it occurs in disturbed areas, and is a common agricultural weed. A native to North America, *A. artemisiifolia* has spread to Europe, Asia, South America and Australia (Friedman and Barrett 2008). Ragweed produces around 1.2 billion grains of pollen per individual (Fumanal et al. 2007). During its flowering season in the late summer and early fall, it is the major cause of hayfever (Bassett et al. 1976), and about 10% of people test positive for allergies to *Ambrosia* (Gergen et al. 1987). In Europe, it is both a public health concern and the cause of crop yield losses (Fenesi and Botta-Dukát 2012). Unsurprisingly, given that ragweed is a wind-pollinated obligate outcrosser, Fst values are low (mean FST=0.025, range= −0.019,0.096) (Martin 2011). Phenotypic and neutral marker diversity are high in the invasive range, providing evidence of multiple introductions (Genton et al. 2005).

### Seed collection and preparation

We collected seed from three populations in both the native (Canada and the United States) and introduced (France) ranges (Figure 1), from overlapping latitudinal ranges (North America: 39.6584-44.37683, France: 43.9475-45.66117) (Colautti et al. 2009). All the populations were large, ranging from hundreds to tens of thousands of individuals. We sampled seed from at least 200 plants in each population. Using methods adapted from Willemsen (1975) and Friedman (pers. comm.), we stratified seeds at 4°C for three months in plastic bags filled with silica and distilled water.

**Figure 1.**
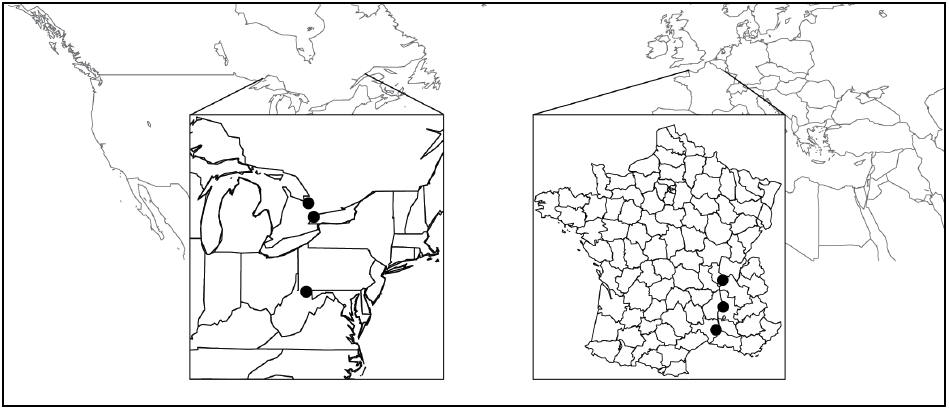
Map of *Ambrosia artemisiifolia* collection sites. Seeds were collected from at least 200 plants in each population in the fall of 2012.

### Parental generation

Beginning in January 2013, we removed seeds from the fridge, and placed them on filter paper in petri dishes. We placed all the petri dishes in the greenhouses and monitored them for germination and signs of dessication. As seeds germinated, we planted them into a 75% Pro-Mix, 20% sand and 5% topsoil soil mix in seedling flats. After four weeks, we transplanted the plants to 4-inch round pots. To keep plants small and accelerate time to flowering, we compressed the growing season. Lights were switched to short days to initiate reproduction. To prevent uncontrolled pollination, we used individual chambers and a purified air delivery system (McGoey et al. 2017). Each plant was placed in a chamber in advance of flowering. Our chambers were composed of plastic bags attached to Styrofoam rings which fit tightly around the pot. The plastic bags were inflated with purified air and individual plants were only removed from the grid for controlled crosses (see McGoey et al. 2017 for more details).

### Crossing design

Breeding designs are critical to partition components of variance in traits (Falconer and Mackay 1996; Lynch and Walsh 1998). We used a nested parental half-sibling design, where we crossed a group of sires (pollen donors) to multiple randomly chosen dams (pollen recipients) (Falconer and Mackay 1996). For each population, we had 50 sires crossed to 3 unique dams each (150 crosses per population) for a total of 900 unique crosses (Conner and Hartl 2003).

### Offspring generation

The offspring generation was grown at the Koffler Scientific Reserve (www.ksr.utoronto.ca; 44.803°N, 79.829°W) during the summer of 2014. As with the parental generation, we stratified seeds and then placed them on petri dishes to germinate. We transplanted seedlings from flat trays four weeks after germination into three blocks. Prior to planting, we removed all vegetation and tilled the soil. At the end of June, seedlings were transplanted over three days into their plots. Each plot had 64 plants each in an 8×8 configuration. Plants were 10 cm apart and plots were 20 cm apart. To promote establishment, we supplemented with water and removed interspecific competitors within the plots for three weeks after transplantation. We also removed interspecific competition adjacent to the plots to prevent shading.

We measured early height (at two weeks), final height, final number of branches, and date of first flower. Most ragweed plants are monoecious (Bassett and Crompton 1975), and so we measured proxies of both male and female fitness. We used the total inflorescence length, which is correlated with pollen production (Fumanal et al. 2007), as a proxy for male reproductive effort. For female reproductive output, we used seed mass which is highly correlated to seed number (r^2^=0.96, p<0.001)(MacDonald and Kotanen 2010).

### Statistical analyses

All statistical analyses were conducted in R version 3.3.1 (R Development Core Team 2016). Historically, it has been difficult to estimate uncertainty around quantitative genetic parameters (Morrissey et al. 2014), but Bayesian Markov chain Monte Carlo (MCMC) methods offer a feasible solution (Hadfield 2015). Using a Bayesian framework enabled us to include uncertainty when estimating the **G** matrices of each population and then carrying those forward through all subsequent analyses and comparisons among populations (Aguirre et al. 2014; Teplitsky et al. 2014).

Estimates of variance components will be constricted to values greater than zero (Walter et al. 2018), which poses challenges for traditional statistical significance testing. To assess the significance of our estimates, we generated a null distribution based on permuting phenotypes within populations. We first created 1000 randomized datasets for each population, where trait values were sampled without replacement and randomly assigned to sires and dams. We used these null expectations in both subsequent univariate and multivariate analyses to assess the significance of estimated variance components—in essence, asking whether the observed phenotypic similarity among related individuals in our experiment was greater than would be expected by random chance.

#### Univariate analyses

To determine if phenotypic traits differed significantly among continents and populations we used models fit with the *MCMCglmm* package in R (Hadfield 2015). We used nested mixed models where block and either population or continent were the fixed factors. Sire and dam were random effects, with dam nested within sire and sire nested within population. We considered a trait mean to be significantly divergent between continents or populations if the HPD intervals for 10000 iterations did not overlap.

We calculated heritabilities from variance components estimated using the *MCMCglmm* package. We estimated the variance among sires (V_S_), among dams within sires (V_D_) and the residual variance (V_E_) (see Supplementary table 1). Since we used a half-sibling breeding design, additive genetic variance (V_A_) was four times the variation in half-sibling families within a population (V_S_). Heritability values in Supplementary table 1 are narrow-sense estimates of V_A_/V_P_. To assess whether our observed heritabilities were greater than expected by chance, we compared them to the 95% HPD intervals of the randomized heritability estimates.

#### Estimation of **G** matrices

We also used the *MCMCglmm* package to generate **G** matrix estimates for the six traits. Block was treated as a fixed effect and dam was nested within sire; we estimated **G** matrices separately for each population. For some of the subsequent analyses, we removed the fitness proxy traits, or treated them separately (noted explicitly below). We present results based on unstandardized data; parallel analyses using traits standardized by the standard deviation (Hansen and Houle 2008) are presented in the supplementary materials/appendix.

#### Comparison of **G** matrices

Researchers have used many different methods to compare **G** matrices (see (Calsbeek and Goodnight 2009; Roff et al. 2012; Aguirre et al. 2014). We sought methods that were biologically interpretable, mathematically tractable, and where statistically uncertainty could be estimated. The biological implications of some metrics can be difficult to ascertain, so here we emphasize methods with clear links to for the evolution of the populations (Aguirre et al. 2014).

To examine overall differences in matrix attributes (i.e. size, shape, orientation) we used Krzanowski’s common subspace analysis and the fourth order genetic covariance tensor. To examine the implications of **G** matrix divergence for the evolutionary trajectories of the populations, we used random skewers, solving the breeder’s equation, and the R metric which predicts evolution with and without covariances (Agrawal and Stinchcombe 2009). Specific details are outlined below.

##### Krzanowski’s common subspace analysis

Some parts of multivariate trait space will have genetic variance, while others will not. We can examine whether the subspaces with the most genetic variation are similar for multiple populations using the Krzanowski subspace analysis (Krzanowski 1979). To find the subspace of most similarity among *p* populations (t =1, …, 6 (in our case)) we can use the equation:

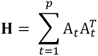

where matrix transposition is indicated by the superscript T and the subset k_t_ of the eigenvectors of G_t_ are contained in A_t_ (Aguirre et al. 2014). The number of eigenvectors included in the summary matrix **H** is half the total number of traits that were examined (Aguirre et al. 2014; Puentes et al. 2016). Any eigenvalues of **H** that are less than *p* indicate that the directions of genetic variation described by that eigenvector differ among populations. In contrast, eigenvalues equal to *p* indicate common sub-spaces—i.e., directions of genetic variation that can be described by the same eigenvectors (Aguirre et al. 2014). The advantage of the Krzanowski method is its clear bounded statistic, which ranges from zero (most divergent) to *p* (most similar) (Blows et al. 2004). Although this method is restricted to examining the subspaces of **G** with the most variation, they are the subspaces that will bias responses to selection and therefore are the most relevant to future adaptation (Aguirre et al. 2014). We conducted the Krzanowski subspace analysis both using all our traits (6 traits, 3 eigenvectors) and while excluding fitness traits (4 traits, 2 eigenvectors). To test for significance, we compared results with those from the randomizations of phenotypes to sires and dams. To consider subspaces significantly diverged, an **H** value had to be lower than *p*, and it had to be lower than the 95% HPD interval calculated from the null **G** matrices.

##### Genetic covariance tensor

The tensor method examines the differences between multiple matrices by using eigenanalyses (Hine et al. 2009). We briefly describe our execution of this analysis; interested readers can refer to Hine et al. (2009) and Aguirre et al. (2014) for more information on the advantages of this approach and details of its implementation. A tensor of the 4^th^ order is required since matrices (2^nd^ order) are being compared (Σ_*ijkl*_ = *cov*(*G*_*ij*_, *G*_*kl*_)). The tensor can be represented as the matrix **S**, containing variances and covariances of all the elements in the original **G** matrices (Hine et al. 2009; Aguirre et al. 2014). Analogously to decomposing a matrix into eigenvalues and eigenvectors, the 4^th^ order tensor can be decomposed into eigenvalues and second-order eigentensors which indicate how the **G** matrices have diverged from one another (Hine et al. 2009; Aguirre et al. 2014). Eigentensors with larger eigenvalues describe axes of variation in the genetic variances and covariances among populations.

The tensor method has the advantages of being compatible with various experimental designs and encompassing all the variation among matrices (Aguirre et al. 2014). Unlike some other methods that may underscore differences that do not easily relate back to the original traits, the 4^th^ order tensor allows for the identification of specific trait combinations that have different variances in the study populations (Aguirre et al. 2014). We used the tensor method on both unstandardized and standardized (see supplementary materials) **G** matrices containing all six traits, and just the four phenotypic traits, using code modified from Aguirre et al. (2014) and Puentes et al. (2016). To determine whether an eigentensor described significant variation among populations we again took advantage of the Bayesian framework (Aguirre et al. 2014). To test whether variation among populations was larger than what could be expected by random sampling error, we compared the posterior distribution of each eigenvalue to the distributions generated from the randomized (null expectation) populations (Aguirre et al. 2014; Careau et al. 2015; Walter et al. 2018). We considered an eigentensor to encompass biologically meaningful variation among populations if the variance it explained was higher than the 95% HPD values calculated from the randomized **G** matrices (null expectations).

##### Random skewers

A primary motivation for comparing **G** matrices is to determine whether the evolutionary trajectories of populations will diverge, and the random skewers approach allows for an investigation of this question (Roff et al. 2012). We can ascertain the collinearity of the responses of two matrices to a series of randomly generated selection vectors (Cheverud and Marroig 2007). Since we had six populations, we did a series of one by one comparisons (i.e. each population was compared to each other population). We used two approaches (described below) to test for differences in the responses of our populations to random skewers.

*Cheverud’s Approach: Angle between response vectors*

For each posterior sample of a **G** matrix, we generated 1000 random vectors, and calculated the multivariate response to selection for each population. For every pair of populations, we then calculated the vector correlation of the responses to selection, and the mean angle between these response vectors. We repeated this for every **G** matrix estimate in our posterior sample (i.e., treating each posterior sample as if it were our only estimate of **G**).

*Aguirre’s Modification: Differences in available genetic variance*

Aguirre and colleagues developed a method based on random skewers that compares the magnitude of genetic variances of multiple populations (Aguirre et al. 2014). In this approach, every randomly generated selection vector is projected through all the iterations of each **G** matrix to estimate genetic variance in the direction of the skewer (cf. Lin and Allaire 1977). For each vector, populations are considered to differ if their each HPD-intervals do not overlap—i.e., they differ in their genetic variance in the direction of the skewer.

As with the first method, 1000 random selection vectors were generated. Each random selection vector was projected through every MCMC iteration of every **G** matrix to estimate genetic variance in the direction of the skewer (Lin and Allaire 1977; Aguirre et al. 2014); consequently for each of the 1000 random vectors, we had posterior distributions of the genetic variance. For each pair of populations, we determined if the highest posterior densities overlapped and collated the cases where they did not.

##### Solving the breeder’s equation

A potential disadvantage of the random skewers method is that the many of the selection skewers that are randomly generated will represent adaptive landscapes that the populations will never confront. The breeder’s equation method uses selection estimates taken empirically. The trait responses of a population facing a realistic selection scenario can then be generated (Lynch and Walsh 1998). Solving the breeder’s equation with a single selection vector can be used to compare **G** matrices: do the matrices lead to significant differences in response to an observed pattern of selection (Stinchcombe et al. 2009) ? Significant differences between populations can be assessed by examining overlaps between the 95% HPD intervals.

We used our field data to estimate directional selection on the four non-fitness traits. In brief, we obtained estimates of Beta by doing a multiple regression of each fitness metric on the phenotypic traits using sire means. We used global sire means to get common representation of the overall pattern of directional selection experienced by our field experimental population (estimating selection for each population separately would make it impossible to distinguish whether differences in observed responses were due to **G** or the selection estimates). We ran separate analyses for male and female fitness as well as a composite of the two (estimated as total inflorescence size, seed mass and the first principal component of a PCA including the two).

##### R values: predicting evolution with and without covariances

Covariances between traits can have important impacts on evolutionary trajectories by either constraining or accelerating adaptation. Agrawal and Stinchcombe (2009) developed a summary statistic to quantify the effect of covariances on the adaptive potential of populations. To implement this method, we calculated the rate of adaptation given our observed **G** matrices and constructed **G** matrices where there are no correlations between traits (i.e. all off-diagonals are changed to zero). The “R value” is the ratio of the two. When R>1, adaptation is accelerated by covariances between traits, when it is <1 is slowed down by genetic correlations.

As with the Breeder’s equation method, the R metric requires selection gradients. We ran separate analyses for male and female fitness, as well as a composite of the two to calculate selection on our four non-fitness traits. To compare populations, we again took advantage of the Bayesian framework and the 10, 000 estimates of each **G** matrix. We compared the R values to determine if the HPD intervals overlapped for each 1×1 comparison of populations.

## Results

### Univariate comparisons

Flowering time varied significantly among populations, while the other traits did not (see Figure 2). Heritability estimates were higher than expected from sampling error for almost all traits and populations (see Figure 3). In some cases, estimates of heritability exceeded one, which has been known to happen due to sampling error (Hill and Thompson 1978).

**Figure 2.**
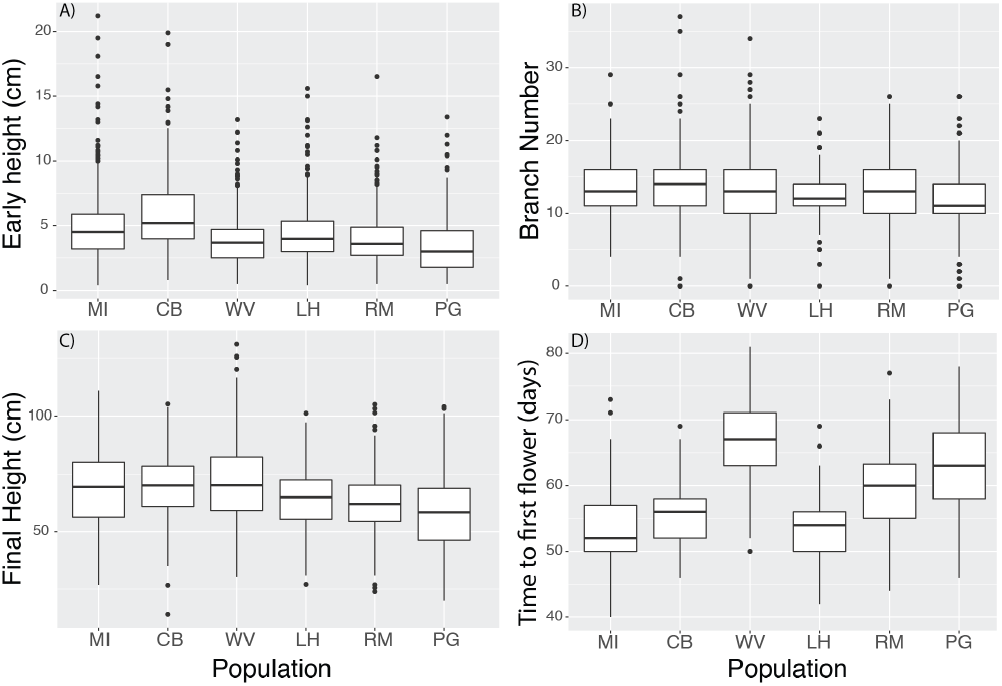
Phenotypic traits of six populations (three native populations from north to south followed by three introduced populations form north to south). Early height (A), branch number (B), final height (C) and flowering time (D).

**Figure 3.**
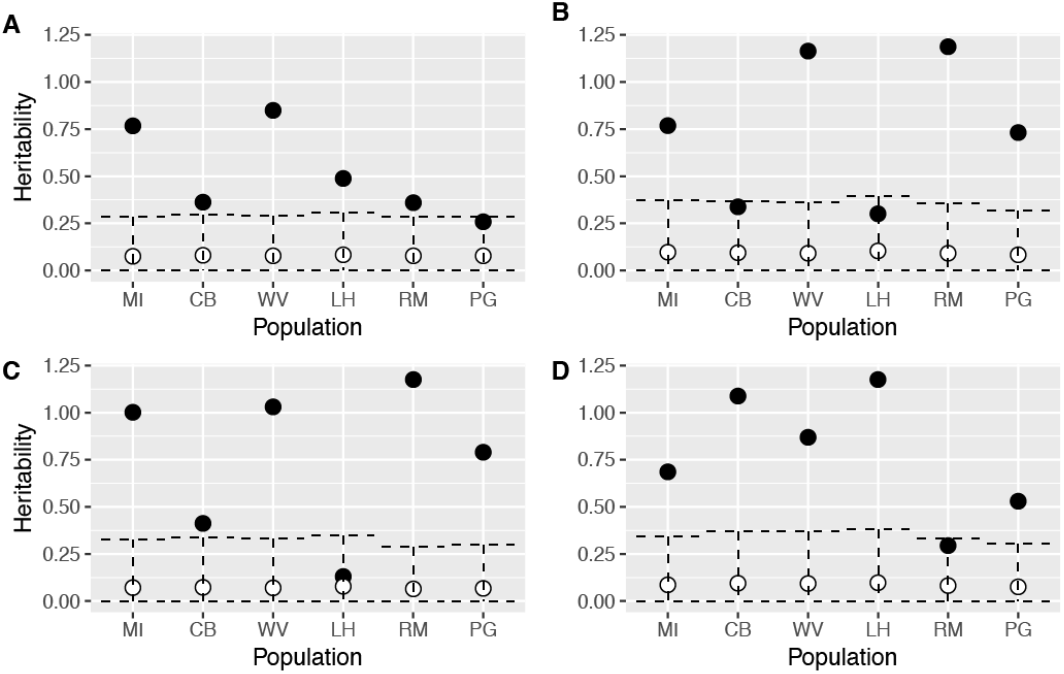
Heritability estimates of early height (A), branch number (B), final height (C) and flowering time(D). Mean posterior estimates are shown in black (circles) and randomized mean estimates are white circles with the 95% intervals shown as dashed lines.

### G matrix comparisons

#### Krzanowski’s common subspace analysis

For our analysis of unstandardized traits, we did not see significant divergences among subspaces among populations based on our criteria (Figure 4). This was true both when all six traits were included, and when we excluded the fitness traits. In other words, the sub-space containing the majority of genetic variation for the six populations was common: there is no evidence of population divergence in the multivariate space described by the leading principal components.

**Figure 4.**
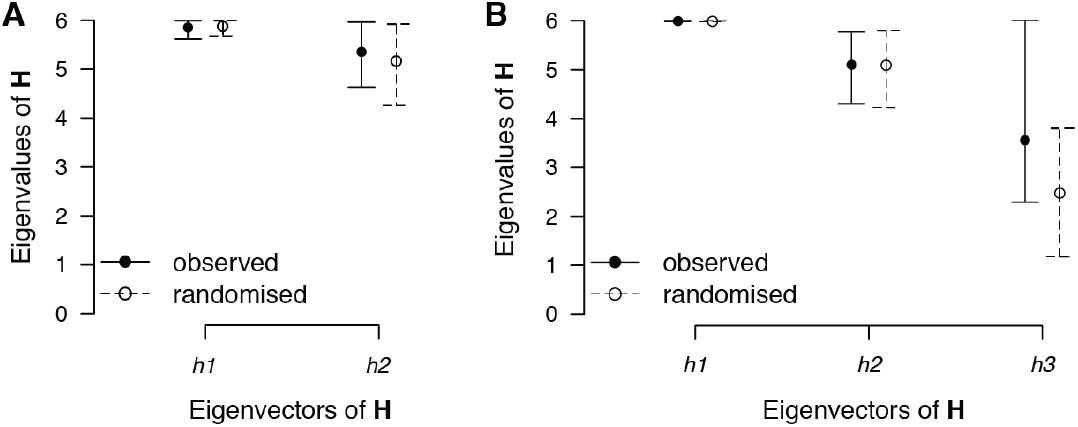
Results from Krzanowski analysis using only the four phenotypic traits (early height, final height, branch number and flowering time) (**A**) and including estimates for male and female fitness (**B**). The eigenvalues (mean and 95 % HPD interval) of each of the first two eigenvectors of H are shown for the observed (closed circle, solid lines) and randomized (open circles, dashed lines)

#### 4^th^ order genetic covariance tensor

For the tensor analysis using all six traits and six populations, we found one significant eigentensor (Figure 5A)—i.e., one direction describing variation and covariation among the six **G** matrices. The first eigentensor described the vast majority of variation (99.5%) among the **G** matrices but there was very large uncertainty in this dimension. The West Virginian population was the most divergent in terms of each populations’ contribution to the first eigentensor. The first eigenvector of the first eigentensor (e11) accounted for 86% of the variation in this eigentensor. The West Virginian population was also divergent in the posterior mean for the genetic variance along the direction of ***e***11 (Figure 5B)). Inflorescence size (our proxy for male fitness) contributed the most to the eigentensor (see supplementary table).

**Figure 5.**
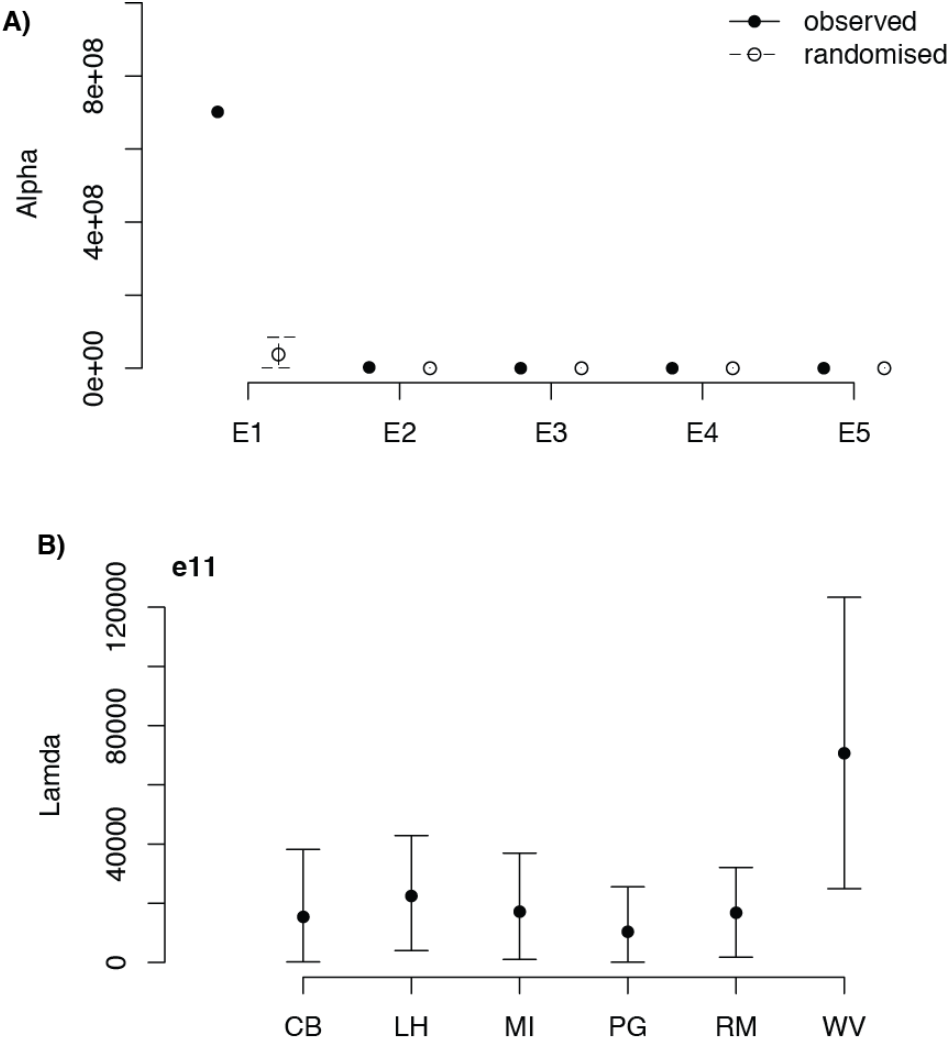
Results of tensor analysis of G matrices for six *Ambrosia artemisiifolia* populations. A) Eigenvalues of eigentensors for posterior mean **S** (the covariance matrix representing the fourth-order covariance tensor). The amount of variance (α) accounted for by each eigentensor is shown for standaredized **G** matrices of the six observed (solid line and circle) and randomized (dashed line and open circle) populations. The error bars are the 95% HPD intervals generated using 10000 MCMC iterations. B) Coordinates of posterior mean for six unstandardized *A. artemisiifolia* **G** matrices in the space of the first eigentensor (E1). Error bars are the 95% HPD intervals for 10000 MCMC samples.

#### Random skewers

We saw mean angles of intermediate value for our one-by-one population comparisons (θ ranging from 0.45 to 0.65 – see supplementary table). For the Aguirre et al. method, we found that when there was divergence between two populations, one of the populations was almost always West Virginia (557 of 563 vectors). Our random skewers results consistently show that divergences between continents were not larger than for populations within continents.

#### Solving the Breeder’s equation

Estimated selection gradients differed in magnitude but not direction depending on the fitness metric that we used (female fitness, male female or a composite of the two). Results from solving the Breeder’s equation were consistent across fitness metrics, so we only present those using composite fitness here (Figure 6). We examined the results of pair-wise comparisons for our six populations for responses in our four non-fitness traits. For unstandardized data, all three size traits (early height, branch number and final height) showed differences in at least one pairwise comparison. As with the random skewers methods, there were not more divergent pairs between continents compared to within continents. In all cases where we saw a difference between populations, one of the populations involved was West Virginia. While these data do not suggest significant differences among populations in the likely response to selection, we did predict significant evolutionary responses in early height, branch number, and final height in 15 out of 18 possible cases. These data indicate that while we predict significant evolutionary responses (i.e., the strength of covariances do not make the predicted response to selection indistinguishable from zero), there is no heterogeneity among our predictions based on populations or continent of origin.

**Figure 6.**
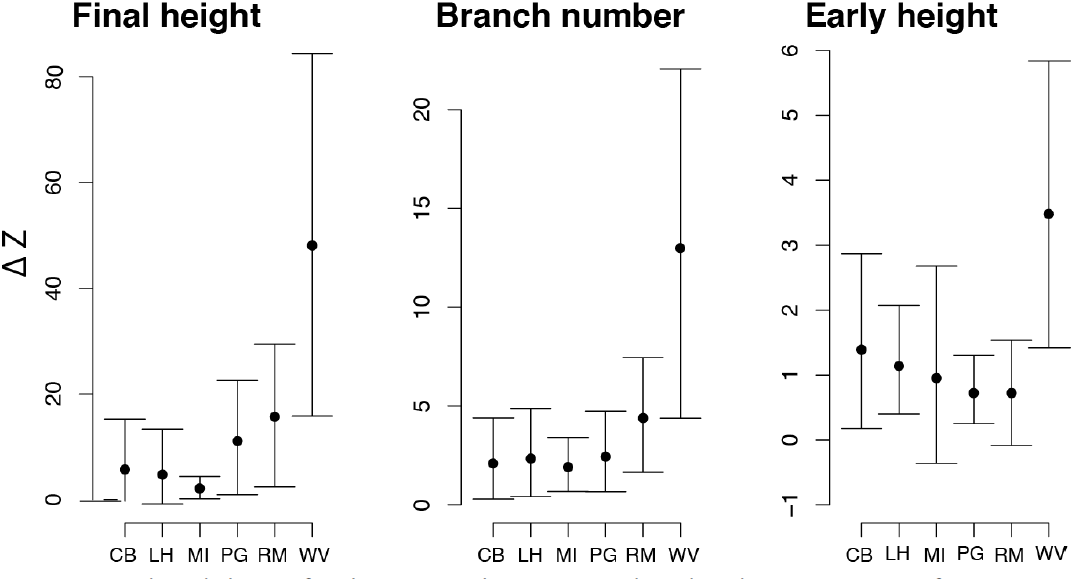
Predicted change for three traits that were predicted to demonstrate significant responses to selection.

#### R values: predicting evolution with and without covariances

As with the decomposition of the breeder’s equation, we generated betas for each of our fitness metrics (female, male and composite). The results did not qualitatively differ, we present those for composite fitness here. To account for uncertainty, we included the 5, 95 percentile as error bars on our plots (see Figure 7). Only West Virginia did not overlap with 1, indicating a significant impact of covariances on its evolutionary trajectory given the same selection scenario. In this case, the average R value was greater than one indicating that covariances would accelerate adaptation.

**Figure 7.**
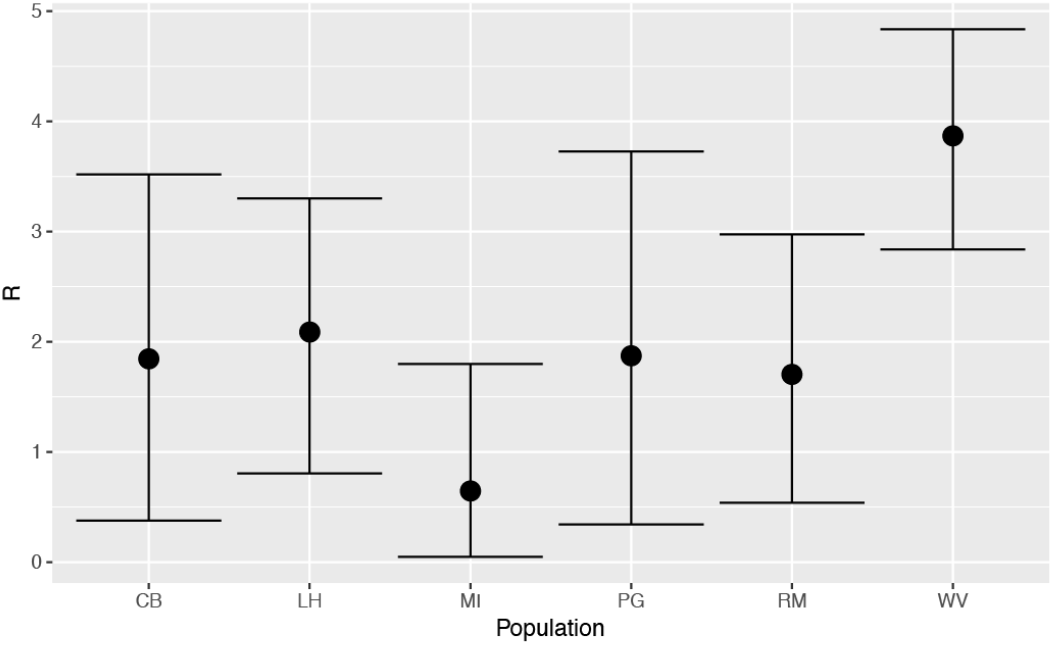
R metric for three native (CB, MI and WV) and three introduced (LH, PG, RM) populations. Values greater than 1 indicate that evolution would be accelerated by genetic correlations and values less than 1 indicate they would be constrained by genetic correlations.

## Discussion

Invasive species are an important component of anthropogenic global change (Simberloff 2014). Invasion genetics examines the importance of genetic factors in determining the trajectory that an invasion will take (Barrett 2015). Most traits that will be important for a response to selection in new habitats will be quantitative (Dlugosch and Parker 2008a; Estoup et al. 2016), which has led to numerous calls for invasion research from a quantitative genetics perspective (Bacigalupe 2009; Lawson Handley et al. 2011) and direct comparisons of additive genetic variance between native and introduced populations (Barrett 2015). Here, we used a common garden experiment paired with multivariate, Bayesian analyses of additive genetic (co)variation to compare the quantitative genetic architecture for native and introduced ragweed populations. While we found some differences in phenotypic traits and their genetic variances, the dominant picture that emerges is that **G** matrices of introduced populations were not significantly or homogeneously diverged from native populations of ragweed. We found that introduced populations did not have lower additive genetic variance or diminished adaptive capacity when compared to native populations. Below we discuss the implications of our results for understanding ragweed’s invasion in particular, and more generally the stability of **G** through space and time.

### Quantitative variation and ragweed invasion

Invasive species represent a major global economic and ecological concern (Pimentel et al. 2005). The field of invasion biology originally emerged from community ecology, and emphasized ecological indicators over evolutionary aspects of introduced populations (Davis 2010). Treating invasive species as static entities may lead to poor predictions on how invasions will proceed (Whitney and Gabler 2008) since evolutionary change occurs on ecological timescales (Thompson 1998). Understanding the role that evolutionary factors such as genetic diversity play in the invasion process is important to our ability to be able to assess and contain invasions (Sakai et al. 2001).

Like many weedy plants, common ragweed has benefited immensely from anthropogenic changes to natural landscapes (Bassett and Crompton 1975; Lavoie et al. 2007). Ragweed is thought to be native to the plains of North America but has spread across the globe. Humans are implicated in every step of this process, from physically transporting it across oceans as a grain contaminant, to constructing roads, to providing consistent disturbances which allow ragweed (an otherwise poor competitor) to persist (Chauvel et al. 2006; Kiss and Béres 2006; Lavoie et al. 2007; MacKay and Kotanen 2008). Recent anthropogenic climate change has extended the growing season for ragweed (Ziska et al. 2011). The consequences of ragweed invasion in Europe are multipronged, including impacts on human health, agricultural productivity and ecological integrity (Chauvel et al. 2006; Buttenschøn et al. 2010). It has been highlighted as a weed of particular concern, with much effort devoted to research on its spread and eradication effort (Pinke et al. 2011).

Ragweed appears to have evolved rapidly in its introduced range, and our results suggest that it has ample quantitative genetic variation for adaptation in traits that could allow further expansion of its range and abundance in Europe. Clines in flowering time and reproductive biomass, and a high Qst (vs Fst) value for reproductive allocation suggest that ragweed has locally adapted across Europe (Hodgins and Rieseberg 2011; Chun et al. 2011). Our results illustrate that the combination of introduction, founder events, and recent adaptation has not reduced quantitative genetic variation relative to source populations: Introduced populations had neither lower heritabilities nor divergent G matrices. Like many other studies, the uncertainty around our G estimates was large, which is a potential qualifier on our conclusion that there is minimal divergence in the **G** matrices between populations (Puentes et al. 2016). The quantitative genetic variation we found is especially concerning given that ragweed also has several characteristics recognized as advantageous for invasion. Ragweed have short generation times, small propagule size and high propagule pressure, all of which will facilitate its spread (Novak 2007; Whitney and Gabler 2008; Dormontt et al. 2010). Outcrossing introduced plants with substantial genetic variation have the capacity to rapidly adapt to their new circumstances (Colautti and Barrett 2013). The traits we focused on are likely critical to the ability of ragweed to continue its range expansion in Europe. Both native and invasive ranges appear to be restricted by phenology (Chapman et al. 2013). Genetic variation for size and flowering time are critical to the ability of invasive species to establish and spread (Colautti and Barrett 2013). The spread of ragweed has been facilitated by railroads and highways, which act as both corridors and habitat (Kiss and Béres 2006; Lavoie et al. 2007). Together, the interconnectedness of Europe, and the high levels of genetic variation already on the continent could accelerate the spread of ragweed into new areas. Eradication efforts of ragweed population must take into account the likelihood of adaptation in response to any interventions and should never treat invasive populations as static.

### Divergences between **G** matrices and their implications

Evolutionary biologists have long held an interest in the stability of **G** over space and time, since the ability to predict evolutionary trajectories are contingent on **G** matrix consistency (Arnold et al. 2008). **G** matrices will be impacted by mutation, selection, drift, recombination and migration (Arnold et al. 2008). The complexities of all these forces interacting have meant that theoretical predictions for how G will change over time have been intractable and the dynamics of G must be studied empirically (Turelli 1988; Revell 2007). There have been several empirical and simulation studies on the stability of **G**, but results are equivocal, and the difficulty in rigorously estimating one **G** matrix, let alone multiple **G** matrices, has meant that we do not yet have a clear picture of how **G** varies in space and time (Arnold et al. 2008; Aguirre et al. 2014; Delahaie et al. 2017). The advent of statistical methods that allow for rigorous comparison of multiple **G** matrices—while accounting for uncertainty in each--has increased the impetus and utility of more empirical research on **G** matrix variability (Delahaie et al. 2017). Despite their importance, studies of **G** matrix variation remain rare, especially for non-model organisms (Cano et al. 2004; Delahaie et al. 2017) and spatial variability is even less well-explored than changes through time (Puentes et al. 2016).

Introduced populations could face two main forces that could shift **G** when compared to native populations. First, a bottleneck could cause a shift in the genetic architecture (Whitlock et al. 2002). Second, the populations could face strong selection which could alter **G** (Arnold et al. 2008). The invasion of ragweed into France has been characterized by multiple introductions and admixture (Genton et al. 2005). Molecular markers show an equivalent or greater diversity in the introduced range, when compared to the native range (Genton et al. 2005). However, the absence of a bottleneck detected from neutral makers does not mean there could not be shifts in quantitative genetic architecture: Neutral markers are not useful as proxies for quantitative genetic variation (Reed and Frankham 2001; Mittell et al. 2015). For example, Eroukhmanoff and Svensson (2011) investigated differences in the **G** matrices of two ecotypes of aquatic isopods. In two different lakes, the isopods have colonized a new habitat in the last few decades. While Eroukmanoff and Svensson (2011) found no difference in neutral genetic variation, additive genetic variance decreased by nearly 50% (Eroukhmanoff and Svensson 2011). Likewise, we cannot use neutral markers to assess adaptive potential. In their study of *Hypericum canariense*, Dlugosch and Parker (2008b) found rapid adaptation of important life history traits in invasive populations, despite large bottlenecks and low molecular genetic diversity.

Our findings, along with past studies (Hodgins and Rieseberg 2011), reveal genetic differentiation for mean values of quantitative traits in ragweed’s introduced range, consistent with divergent directional selection since colonization. There have also been multiple introductions from different source populations, potentially causing shifts in **G** due to waves of migration. Despite the different evolutionary forces introduced ragweed populations have faced, their **G** matrices have not substantially diverged from those of native populations. We used a variety of methods to assess the magnitude of **G** matrices differences for native and introduced ragweed populations. Overall the **G** matrices are largely stable across geography, consistent with studies on other taxa that have also found similarity in **G** across conspecific populations. While we did find some moderate differences between **G** matrices, most differences seem to be driven by the West Virginian population, which was highlighted by several of the analyses as a divergent population. Differences were not more apparent between populations from different continents than those from the same range. When confronted with the same selection scenario, responses of introduced populations would not be more different from native populations than from each other. There are too few studies of G matrix variability among populations for broad patterns to emerge, but past authors have argued that G-matrices are stable across geography (Arnold et al. 2008; Delahaie et al. 2017).

### Conclusion

It is increasingly appreciated that evolutionary factors are important in the invasion process, and that there is value in approaching the study of invasive species from a quantitative genetics perspective. Data on the adaptive potential of wild populations are scarce (Delahaie et al. 2017), but are necessary to understanding evolution in natural environments.

Here, we have used a multivariate, Bayesian approach and found that introduced *A. artemisiifolia* populations are not limited in their adaptive potential when compared to native populations. Importantly, the availability of additive genetic variance seen here indicates that ragweed will be able to respond to selection pressures in the introduced range, whether from novel selection, global change, or eradication efforts. Combined with its annual life history and prolific production of seeds, ragweed is primed to adapt rapidly to selection pressures that arise in its introduced range.

